# *In vitro* activity of the new β-lactamase inhibitors relebactam and vaborbactam in combination with β-lactams against *Mycobacterium abscessus* complex clinical isolates

**DOI:** 10.1101/499830

**Authors:** Amit Kaushik, Nicole C. Ammerman, Jin Lee, Olumide Martins, Barry N. Kreiswirth, Gyanu Lamichhane, Nicole M. Parrish, Eric L. Nuermberger

**Affiliations:** Center for Tuberculosis Research, Johns Hopkins University School of Medicine, Baltimore, Maryland, USA; Public Health Research Institute Tuberculosis Center, New Jersey Medical School - Rutgers, The State University of New Jersey, Newark, New Jersey, USA; Department of Pathology, Johns Hopkins University School of Medicine, Baltimore, Maryland, USA

**Keywords:** β-lactamase inhibitors, β-lactams, relebactam, vaborbactam, carbapenems, cephalosporins, *Mycobacterium abscessus*

## Abstract

Pulmonary disease due to infection with *Mycobacterium abscessus* complex (MABC) is notoriously difficult to treat, in large part due to MABC’s intrinsic resistance to most antibiotics, including β-lactams. MABC organisms express a broad-spectrum β-lactamase that is resistant to traditional β-lactam-based β-lactamase inhibitors but inhibited by a newer non-β-lactam-based β-lactamase inhibitor, avibactam. Consequently, the susceptibility of MABC to some β-lactams is increased in the presence of avibactam. Therefore, we hypothesized that two new non-β-lactam-based β-lactamase inhibitors, relebactam and vaborbactam, would also increase susceptibility of MABC to β-lactams. The objective of the present study was to evaluate the *in vitro* activity of various marketed β-lactams alone and in combination with either relebactam or vaborbactam against multidrug-resistant MABC clinical isolates. Our data demonstrate that both β-lactamase inhibitors significantly improved the anti-MABC activity of many carbapenems (including imipenem and meropenem) and cephalosporins (including cefepime, ceftaroline, and cefuroxime). As a meropenem/vaborbactam combination is now marketed and an imipenem/relebactam combination is currently in phase III trials, these fixed combinations may become the β-lactams of choice for the treatment of MABC infections. Furthermore, given the evolving interest in dual β-lactam regimens, our results identify select cephalosporins, such as cefuroxime, with superior activity in the presence of a β-lactamase inhibitor, deserving of further evaluation in combination with these carbapenem/β-lactamase inhibitor products.

## Introduction

*Mycobacterium abscessus* (or *M. abscessus* subsp. *abscessus*), *M. massiliense* (or *M. abscessus* subsp. *massiliense*), and *M. bolletii* (or *M. abscessus* subsp. *bolletii*) comprise the *M. abscessus* complex (MABC) (1). These rapidly-growing nontuberculous mycobacteria, ubiquitous in the environment, are opportunistic human pathogens associated with a wide range of maladies, from localized skin lesions to systemic disease. Individuals with cystic fibrosis and other forms of bronchiectasis are especially vulnerable to MABC pulmonary disease, an infection that is notoriously difficult to eradicate due in large part to MABC’s broad, intrinsic resistance to most antibiotics, including many anti-mycobacterial drugs (2–4). The paucity of effective treatment regimens has recently gained attention as the prevalence of MABC pulmonary disease is apparently increasing (5–7), justly highlighting the need for additional treatment options.

Like several other pathogenic and nonpathogenic mycobacteria, MABC organisms possess a constitutively expressed, broad-spectrum β-lactamase, BlaMab, which contributes to the intrinsic resistance of MABC to most β-lactam antibiotics (8–12). Several studies have indicated that BlaMab is not significantly inhibited by β-lactam-based β-lactamase inhibitors, namely clavulanate, tazobactam, and sulbactam (9, 13–15). In contrast, the non-β-lactam-based β-lactamase diazabicyclooctane (DBO) inhibitor avibactam does inhibit BlaMab, thereby reducing the minimum inhibitory concentration (MIC) of many β-lactams for MABC, especially carbapenems and cephalosporins, to clinically achievable concentrations (16–20). Avibactam is marketed solely in combination with the cephalosporin ceftazidime (trade name AVYCAZ^®^ in the United States). However, ceftazidime has little or no demonstrable activity against MABC, even in combination with avibactam and against *M. abscessus* strains in which the gene encoding BlaMab has been entirely deleted (8, 9, 18). Thus, the current requirement to co-administer ceftazidime in order to potentiate the activity of other more effective β-lactams with avibactam complicates this treatment strategy for MABC infections, as ceftazidime might only incur risk of adverse effects without perceived benefit.

Relebactam and vaborbactam are two newer non-β-lactam-based β-lactamase inhibitors developed for use with the carbapenems imipenem and meropenem, respectively (21). Whereas relebactam is a DBO β-lactamase inhibitor structurally related to avibactam, vaborbactam is a novel boronic acid-based inhibitor. While neither of these β-lactamase inhibitors are expected to be clinically available as sole formulations, both of the paired carbapenems have activity against MABC. Imipenem alone has good activity and is currently recommended as part of first-line treatments for MABC pulmonary disease (2, 3). Activity of meropenem, while comparatively less than imipenem when used alone, is increased comparable to that of imipenem in the presence of avibactam (8, 16, 18). As the meropenem/vaborbactam combination is already clinically available (trade name VABOMERE™ in the United States), and the imipenem/cilastatin/relebactam combination is currently being evaluated in multiple phase III clinical trials (ClinicalTrials.gov identifiers NCT02493764, NCT03583333, NCT03293485, NCT02452047), we set out to assess the impact of these β-lactamase inhibitors on the anti-MABC activity of a variety of β-lactam drugs. The objective of this study was to evaluate the activity of β-lactams alone and in combination with either relebactam or vaborbactam, against MABC, including multidrug-resistant (MDR) clinical isolates.

## Results

### Impact of culture medium on the *in vitro* growth of MABC clinical isolates

Clinical and Laboratory Standards Institute (CLSI) guidelines recommend the use of cation-adjusted Mueller-Hinton broth (CAMHB) for susceptibility testing of antimicrobials against rapidly-growing mycobacteria, including MABC; for MIC determination, the guidelines state that cultures should be examined after 3 days of incubation, to be extended up to 5 days if growth of the non-drug-containing control sample is insufficient (22). Early in our work, we found that MABC clinical isolates in our collection, isolates resistant to almost all antimicrobials currently used to treat MABC infection (16), grow slowly in CAMHB, and that, on average, MIC values could not be determined until nearly 5 days of incubation (**Fig. S1A**). Such a long incubation period can be problematic when evaluating the activity of some β-lactams due to their innate instability in aqueous media, independent of the presence of β-lactamase enzymes (8, 23–25), which could potentially result in artificially high MIC values. The clinical strains grow better in Middlebrook 7H9 broth supplemented with 10% (v/v) oleic acid-albumin-dextrose-catalase (OADC) enrichment (**Fig. S1B**), a liquid laboratory medium for culturing mycobacteria (26–28). Therefore, Middlebrook 7H9 liquid medium was primarily used in this study.

### Activity of β-lactams with relebactam or vaborbactam against M. *abscessus* ATCC 19977

We first evaluated the impact of relebactam and vaborbactam on the activity of β-lactams against a well-characterized MABC strain, *M. abscessus* ATCC 19977 (29). MICs of drugs representing the four major sub-classes of β-lactams (carbapenems, cephalosporins, monobactams, and penicillins) were determined in the presence and absence of relebactam or vaborbactam at 4 μg/ml (Table 1). Neither of these β-lactamase inhibitors exhibited antimicrobial activity on their own. The MICs of relebactam alone and vaborbactam alone were both >256 μg/ml, and neither drug inhibited bacterial growth in a disk diffusion assay (**Fig. S2**). As expected, without co-exposure to a β-lactamase inhibitor, the strain was able to grow in relatively high concentrations of all β-lactams tested, with imipenem having the lowest MIC at 8 μg/ml. Overall, the MICs of penicillins and the monobactam aztreonam were not affected by either β-lactamase inhibitor, although the MIC of amoxicillin did shift from >256 μg/ml to 32 μg/ml in the presence of relebactam. In contrast, all carbapenems and more than half of the cephalosporins tested exhibited decreased MICs in the presence of either relebactam or vaborbactam. The magnitude of the MIC shift associated with each β-lactamase inhibitor was similar for most of these β-lactams, but for two of the carbapenems (tebipenem and ertapenem) and three of the cephalosporins (cefdinir, cefuroxime, and ceftaroline) the MICs associated with relebactam were one dilution lower than that associated with vaborbactam. Similar activity was also observed in a disk diffusion assay (**Fig. S2**).

**TABLE 1.**
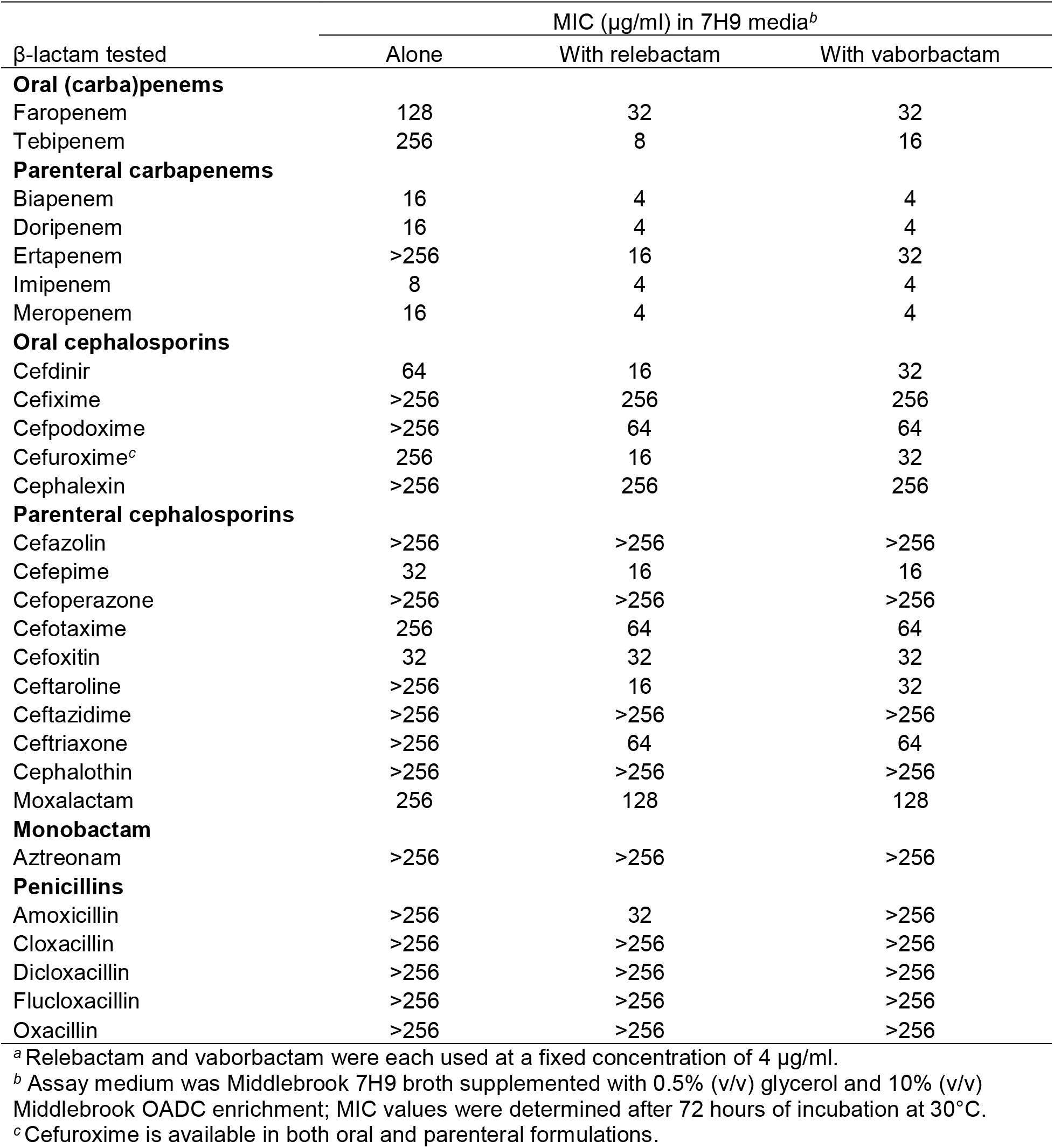
MIC values of β-lactams with and without β-lactamase inhibitors^a^ against *M. abscessus* strain ATCC 19977

### Activity of β-lactams with relebactam or vaborbactam against MABC clinical isolates

The promising antimicrobial activity of the carbapenems and select cephalosporins, namely cefdinir, cefpodoxime, cefuroxime, cefepime, cefotaxime, cefoxitin, ceftaroline, and ceftriaxone, in combination with either relebactam or vaborbactam against the ATCC 19977 strain prompted us to evaluate the activity of these combinations against a collection of 28 MABC clinical isolates with MDR phenotypes (16). The MICs of the carbapenems and cephalosporins against each clinical isolate are presented in Tables S1 and S2, respectively. With the exception of cefoxitin, the MICs of each β-lactam decreased in the presence of either β-lactamase inhibitor (Table 2). As observed with the ATCC 19977 strain, both β-lactamase inhibitors were mostly associated with MIC shifts of similar magnitude, although the MIC_50_, MIC_90_, and MIC range limits associated with relebactam were one or two dilutions lower than those associated with vaborbactam for tebipenem, biapenem, and meropenem. For tebipenem, 25/28 (89%) isolates has lower MIC values with relebactam compared to vaborbactam, and for biapenem and meropenem, 14/28 (50%) and 10/28 (36%) of the isolates, respectively, had lower MICs with relebactam compared to vaborbactam. For the carbapenems, the most striking shifts in MIC distribution were observed with tebipenem and ertapenem (**Fig. 1A,B**), while the lowest MIC values were obtained with imipenem, meropenem, biapenem, and doripenem (**Fig. 1C-F**). The MIC_50_ value for each of these carbapenems was 4-8 μg/ml in the presence of either β-lactamase inhibitor, suggesting that they have similar intrinsic activity in the absence of hydrolysis by BlaMab. For the cephalosporins, the largest shifts in MIC distribution associated with the β-lactamase inhibitors were observed for ceftaroline, ceftriaxone, and cefuroxime (**Fig. 2A-C**), while the shifts were comparatively smaller for cefotaxime and cefepime (**Fig. 2D,E**). The MIC distribution of cefoxitin was not affected by the presence of either relebactam or vaborbactam (**Fig. 2F**). The lowest MICs in association with a β-lactamase inhibitor were observed with cefuroxime, cefepime, cefdinir, and ceftaroline.

**TABLE 2.**
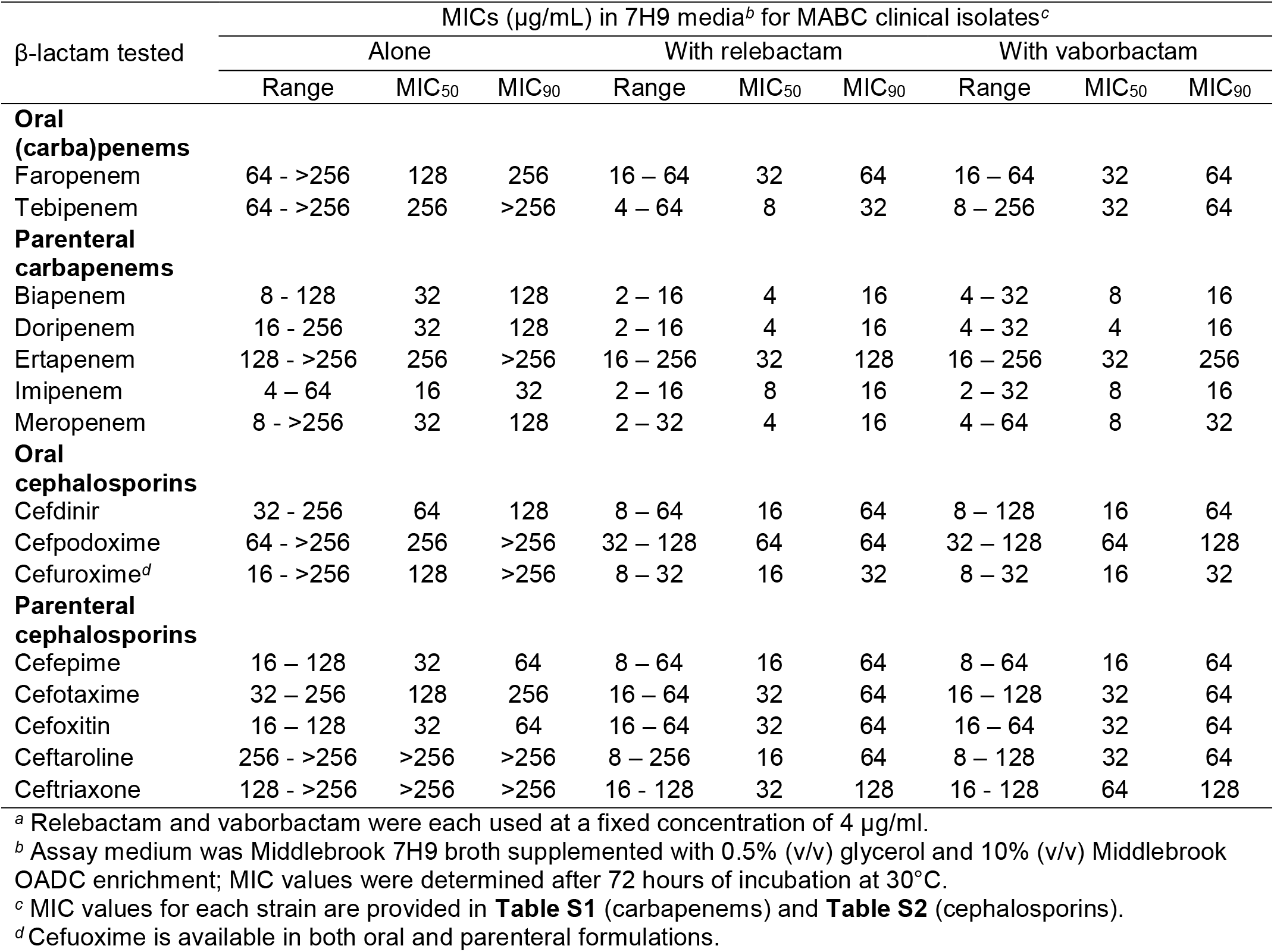
MIC values of β-lactams with and without β-lactamase inhibitors^a^ against 28 MDR MABC clinical isolates

| β-lactam tested | MICs (μg/mL) in 7H9 media<sup>b</sup> for MABC clinical isolates<sup>c</sup> | | | | | | | | |
| --- | --- | --- | --- | --- | --- | --- | --- | --- | --- |
|  | Alone |  |  | With relebactam |  |  | With vaborbactam |  |  |
|  | Range | MIC<sub>50</sub> | MIC<sub>90</sub> | Range | MIC<sub>50</sub> | MIC<sub>90</sub> | Range | MIC<sub>50</sub> | MIC<sub>90</sub> |
| **Oral (carba)penems** |  |  |  |  |  |  |  |  |  |
| Faropenem | 64 - >256 | 128 | 256 | 16 - 64 | 32 | 64 | 16 - 64 | 32 | 64 |
| Tebipenem | 64 - >256 | 256 | >256 | 4 - 64 | 8 | 32 | 8 - 256 | 32 | 64 |
| **Parenteral carbapenems** |  |  |  |  |  |  |  |  |  |
| Biapenem | 8 - 128 | 32 | 128 | 2 - 16 | 4 | 16 | 4 - 32 | 8 | 16 |
| Doripenem | 16 - 256 | 32 | 128 | 2 - 16 | 4 | 16 | 4 - 32 | 4 | 16 |
| Ertapenem | 128 - >256 | 256 | >256 | 16 - 256 | 32 | 128 | 16 - 256 | 32 | 256 |
| Imipenem | 4 - 64 | 16 | 32 | 2 - 16 | 8 | 16 | 2 - 32 | 8 | 16 |
| Meropenem | 8 - >256 | 32 | 128 | 2 - 32 | 4 | 16 | 4 - 64 | 8 | 32 |
| **Oral cephalosporins** |  |  |  |  |  |  |  |  |  |
| Cefdinir | 32 - 256 | 64 | 128 | 8 - 64 | 16 | 64 | 8 - 128 | 16 | 64 |
| Cefpodoxime | 64 - >256 | 256 | >256 | 32 - 128 | 64 | 64 | 32 - 128 | 64 | 128 |
| Cefuroxime<sup>d</sup> | 16 - >256 | 128 | >256 | 8 - 32 | 16 | 32 | 8 - 32 | 16 | 32 |
| **Parenteral cephalosporins** |  |  |  |  |  |  |  |  |  |
| Cefepime | 16 - 128 | 32 | 64 | 8 - 64 | 16 | 64 | 8 - 64 | 16 | 64 |
| Cefotaxime | 32 - 256 | 128 | 256 | 16 - 64 | 32 | 64 | 16 - 128 | 32 | 64 |
| Cefoxitin | 16 - 128 | 32 | 64 | 16 - 64 | 32 | 64 | 16 - 64 | 32 | 64 |
| Ceftaroline | 256 - >256 | >256 | >256 | 8 - 256 | 16 | 64 | 8 - 128 | 32 | 64 |
| Ceftriaxone | 128 - >256 | >256 | >256 | 16 - 128 | 32 | 128 | 16 - 128 | 64 | 128 |

<sup>a</sup> Relebactam and vaborbactam were each used at a fixed concentration of 4 μg/ml.
<sup>b</sup> Assay medium was Middlebrook 7H9 broth supplemented with 0.5% (v/v) glycerol and 10% (v/v) Middlebrook OADC enrichment; MIC values were determined after 72 hours of incubation at 30°C.
<sup>c</sup> MIC values for each strain are provided in **Table S1** (carbapenems) and **Table S2** (cephalosporins).
<sup>d</sup> Cefuroxime is available in both oral and parenteral formulations.

**FIG. 1.**
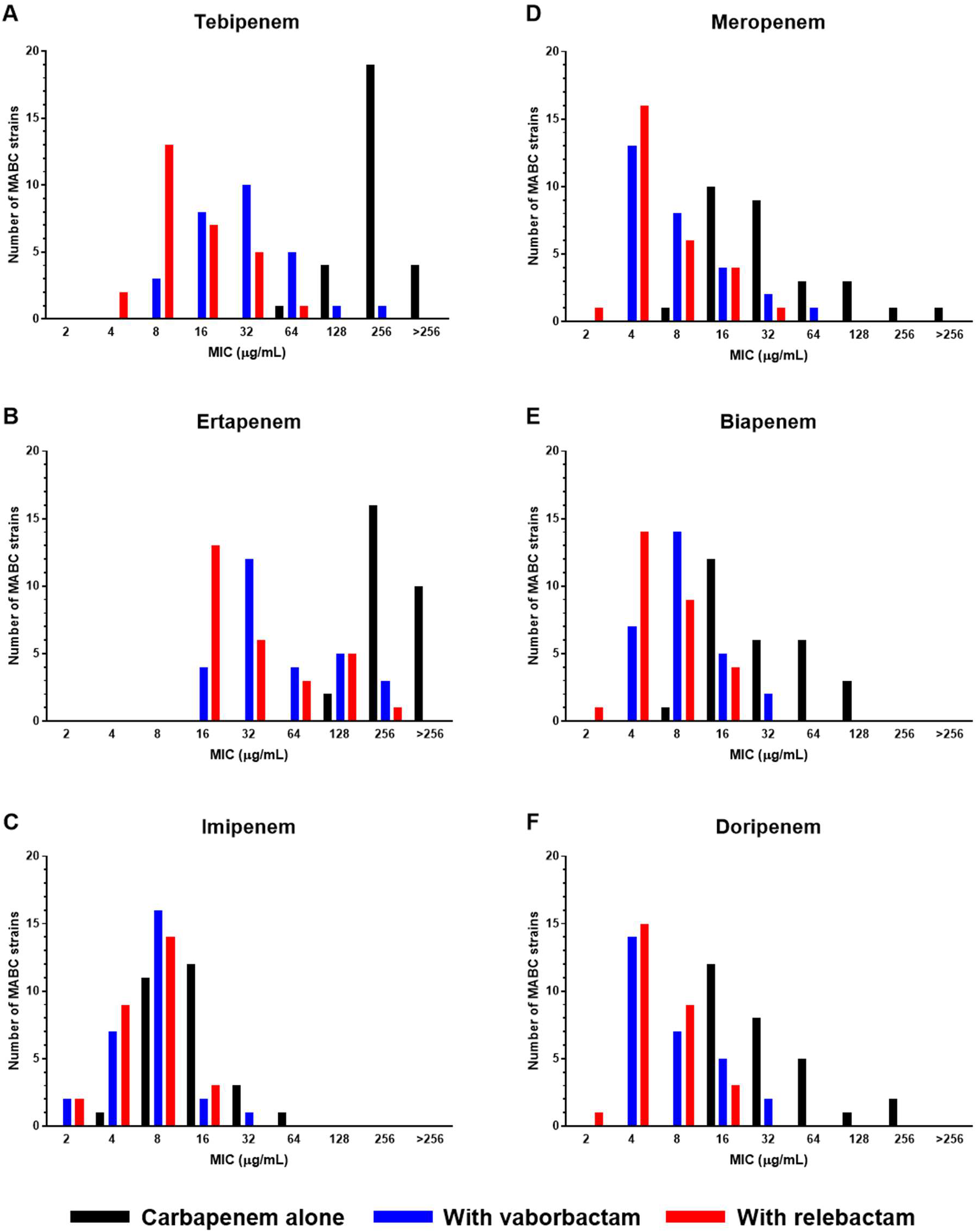
MIC distributions of carbapenems, alone and in combination with 4 μg/ml relebactam or vaborbactam, against 28 MABC clinical isolates.

**FIG. 2.**
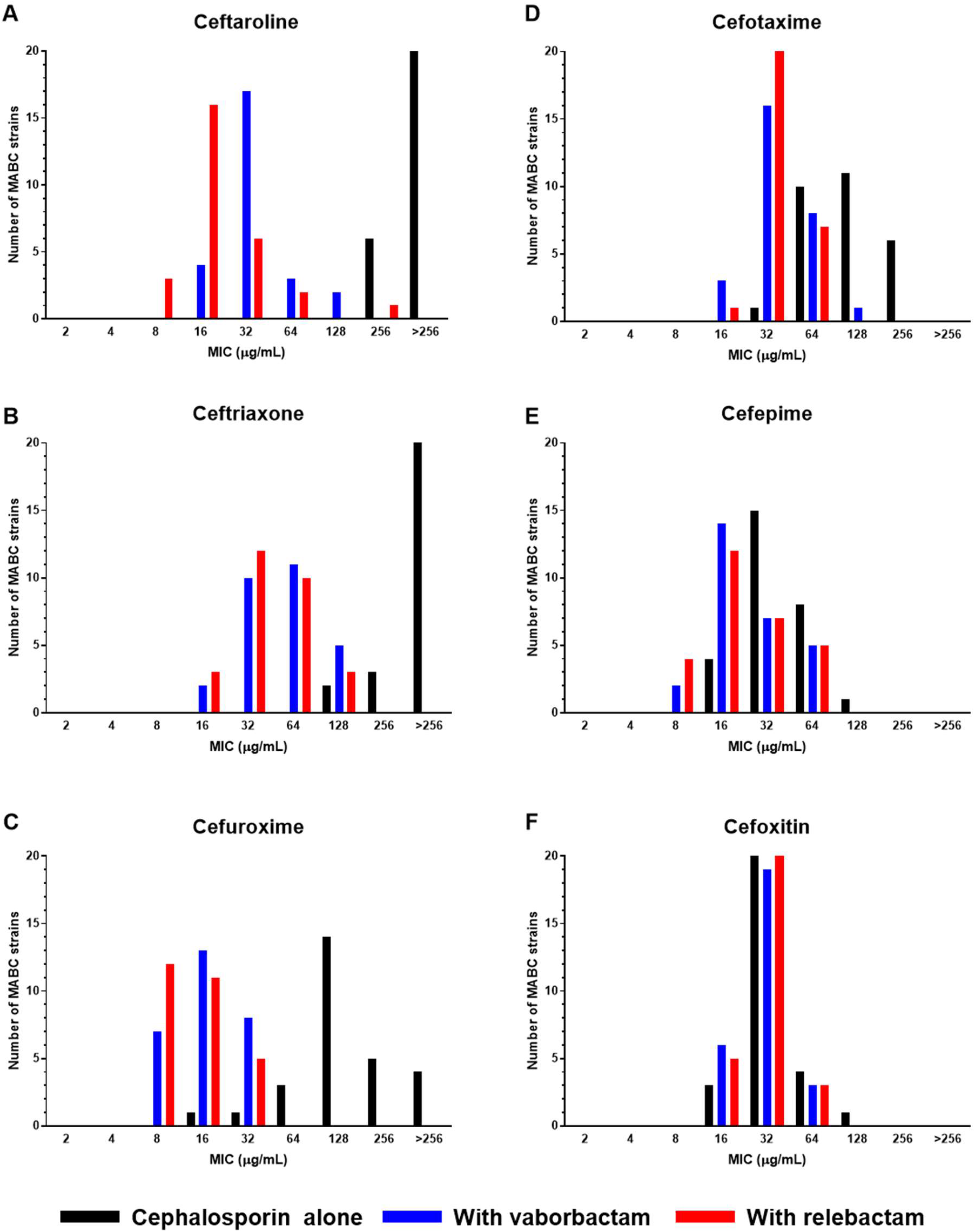
MIC distributions of cephalosporins, alone and in combination with 4 μg/ml relebactam or vaborbactam, against 28 MABC clinical isolates.

For a selection of β-lactams, MIC values with and without the β-lactamase inhibitors were also determined in CAMHB. For the ATCC 19977 strain, the MICs were determined after 3 days of incubation, while the MICs for clinical isolates could not be determined before day 4 or 5 of incubation (**Table S3**). For three of the clinical isolates (strains 2N, 11N, and JHHKB), adequate growth for determining MIC values did not occur within 5 days. With the exception of imipenem and cefdinir, the MIC_50_ values of the tested β-lactams were higher, both the presence and absence of relebactam or vaborbactam (**Table S4**), relative to the MIC_50_ values determined in Middlebrook 7H9 liquid medium (Table 2). The MIC values the ATCC 19977 strain also tended to be higher in CAMHB (**Table S3**).

## Discussion

This study demonstrates that certain β-lactams, namely carbapenems and cephalosporins, exhibit improved activity against MABC in the presence of vaborbactam or relebactam, two new non-β-lactam-based β-lactamase inhibitors. Vaborbactam and relebactam reduced the MIC_50_, MIC_90_, and MIC range limits of meropenem and imipenem, respectively, by one or two dilutions in both 7H9 and CAMHB media. Since meropenem and imipenem MICs against MABC clinical isolates commonly lie at or around the recommended susceptibility breakpoints, these carbapenem/β-lactamase inhibitor combinations can be expected to improve PK/PD target attainment for the carbapenems and increase efficacy over either carbapenem alone. We recently observed similar reductions of carbapenem MICs with avibactam against the same clinical isolates (16). However, as avibactam is currently marketed only with ceftazidime, a cephalosporin with poor intrinsic activity itself against MABC, the carbapenem/β-lactamase inhibitor combinations studied here appear more advantageous by not requiring combination of the carbapenem with ceftazidime/avibactam, which imposes the risk of ceftazidime-related side effects in order to gain the value of Bla_Mab_ inhibition. Further confirmation of these results against a larger number of clinical isolates might promote meropenem/vaborbactam or, pending future regulatory approval, imipenem/relebactam to become the β-lactams of choice against MABC infections.

The significant reductions in MIC values for carbapenems and cephalosporins against MABC in the presence of vaborbactam and relebactam provide indirect evidence that both of these β-lactamase inhibitors inhibit BlaMab, thus revealing the intrinsic antimicrobial activity of the paired β-lactams against their transpeptidase targets in MABC (30). Although carbapenems clearly have greater intrinsic potency than the cephalosporins against the MABC isolates tested here, vaborbactam and relebactam, like avibactam previously (16), also significantly reduced the MICs of a variety of marketed cephalosporins. A key question is whether these β-lactamase inhibitors increase the susceptibility of MABC to the cephalosporins to levels that are clinically relevant based on drug exposures that are achievable in patients.

One way to address this question is to compare the observed MICs in the context of CLSI interpretive categories and MIC breakpoints set for the antimicrobial susceptibility testing of these β-lactams (22, 31). Breakpoints for the interpretive categories of susceptible, intermediate, and resistant are not based solely on the natural MIC distribution, but also on microbiological and pharmacological data, and are “considered to be robust predictors of clinical outcome” (31). Among the drugs tested in this study, recommended breakpoints for rapidly-growing mycobacteria are currently only available for imipenem, meropenem, and cefoxitin (**Table S5**), three β-lactams currently recommended for use for treatment of mycobacterial infections and expected to reach active concentrations in patients (2, 3). However, CLSI-recommended breakpoints for infections caused by other bacterial genera are available for all of the cephalosporins tested in this study (31). Although these cephalosporin breakpoints should not be used to predict susceptibility of MABC for clinical decision-making at this time, they can be considered as surrogate indicators of potential susceptibility to drug exposures that are achievable in patients. As previously observed for avibactam (16), vaborbactam and relebactam did not significantly affect the MICs of cefoxitin (MIC_50_ 32 μg/ml with or without either β-lactamase inhibitor) against our isolates (**Tables 1, 2**), further confirming cefoxitin’s stability in the presence of BlaMab (9), while also calling into question whether selected cephalosporins could be superior to cefoxitin for treatment of MABC if combined with an effective β-lactamase inhibitor. However, the addition of vaborbactam and/or relebactam decreased the MIC_50_ values of cefotaxime, ceftriaxone, cefuroxime and cefepime for MABC to concentrations at or within one dilution of the CLSI susceptible/intermediate breakpoints for these drugs against *Enterobacteriaceae* or anaerobic bacteria (**Table S5**), thus indicating that MABC may be susceptible to clinically achievable concentrations of these cephalosporins in the presence of vaborbactam or relebactam. With MICs consistently one dilution lower than cefoxitin in the presence of vaborbactam or relebactam, cefuroxime is particularly attractive given its narrower spectrum of activity, longer half-life and lower plasma protein binding compared to cefoxitin. Moreover, at a standard dose of 1.5 g every 8 hours, cefuroxime is expected to have higher probability than cefoxitin at a dose of 2 g every 8 hours of achieving adequate T_>MIC_ even when MIC distributions for the two drugs are similar, as shown by Moine *et al.* (32). Further studies are needed to determine whether cefuroxime or an alternative cephalosporin might have superior efficacy compared to the currently recommended agent cefoxitin when combined with a β-lactamase inhibitor.

Recent studies suggest that combining different classes of β-lactams against MABC (30, 33) and other bacteria (34, 35) may be synergistic due to targeting a wider spectrum of enzymes that appear to be uniquely relevant to synthesis of peptidoglycan in MABC (36, 37). For example, Kumar *et al.* recently demonstrated synergy between doripenem and cefdinir against MABC (30). Remarkably, despite its poor activity when tested alone or in combination with avibactam, ceftazidime demonstrates synergy with either imipenem or ceftaroline against MABC, whether avibactam is present or not (38). These studies suggest that a dual-β-lactam combination may be superior to a single β-lactam for treatment of MABC pulmonary disease. In this context, our finding that several marketed cephalosporins exhibit greater activity than ceftazidime in the presence of vaborbactam and relebactam indicates that administering a carbapenem/β-lactamase inhibitor combination together with a more intrinsically active cephalosporin, such as cefuroxime, may have superior activity compared to the same carbapenem combined with ceftazidime/avibactam or another cephalosporin or the same carbapenem/β-lactamase inhibitor alone.

Another interesting finding of our study is that the MICs of the carbapenems and cephalosporins, alone and in combination with relebactam or vaborbactam, against the ATCC 19977 strain were often similar to MICs against the MABC clinical isolates (**Tables S1, S2**). For all of the 15 drugs tested, the MIC for the ATCC 19977 strain fell within one dilution of the MIC_50_ for the clinical isolates (**Tables 1, 2**). Thus, the MDR clinical isolates, which were isolated from patients with cystic fibrosis and therefore likely previously exposed to β-lactam drugs, were still largely vulnerable to those cephalosporins and carbapenems with activity against MABC transpeptidase targets. These findings also suggest that the widely used and well-characterized *M. abscessus* strain ATCC 19977 (29) is a suitable model strain for the evaluation of β-lactam activity, with or without β-lactamase inhibitors, and therefore can serve as a valuable tool for studying the molecular mechanisms of intrinsic susceptibility and resistance of MABC to β-lactams.

CLSI guidelines state that broth MIC testing of rapidly-growing mycobacteria, including MABC, should be performed using CAMHB (22), but we found that our MABC clinical isolates grew poorly in this medium, allowing MIC reading to be done after 4, 5, or even 6 days of incubation (**Fig. S1, Table S3**). Due to concerns that the inherent instability of β-lactams may lead to artificially high MIC values after such prolonged incubation periods (8, 23–28), we relied primarily on Middlebrook 7H9 liquid medium for the broth MIC assays in this study. However, we also evaluated a selection of carbapenems and cephalosporins in CAMHB, and found that, in general, the MICs were indeed higher in CAMHB compared to Middlebrook 7H9 medium (Tables 2, S4). We also observed that MICs against strain ATCC 19977, which grew equivalently in either type of media, tended to be higher in CAMHB, indicating that media-specific factors may influence the anti-MABC activity of β-lactams. There are other reports of higher β-lactam MICs against strain ATCC 19977 in CAMHB compared to in Middlebrook 7H9 liquid medium (16, 18, 19). As CAMHB is recommended for use in clinical microbiology laboratories for antimicrobial susceptibility testing of MABC isolates from patient samples, it will be important to understand the basis of these apparent media-dependent discrepancies in MABC susceptibility to β-lactams. Ultimately, the imperative is to understand which MICs in any given medium correlate with *in vivo* activity. Preclinical animal models suitable for testing the antimicrobial activity of β-lactams against MABC would be useful to address this issue, but the absence of a well-qualified animal model is currently a critical missing link in our ability to translate *in vitro* activity to clinical utility.

In conclusion, the data presented in this study demonstrate that the β-lactamase inhibitors vaborbactam and relebactam markedly improve the anti-MABC activity of many carbapenems and cephalosporins and should improve the clinical utility of some of these β-lactams. Each of these β-lactamase inhibitors is currently formulated with a carbapenem: vaborbactam with meropenem and relebactam with imipenem, and our data suggest that these fixed combinations will be more useful for the treatment of MABC disease than either carbapenem alone, especially in the case of meropenem. Moreover, the combinations of meropenem/vaborbactam and, if it is approved in the future, imipenem/relebactam, would make it unnecessary to combine the carbapenems with the marketed ceftazidime/avibactam combination in order to gain effective β-lactamase inhibition. Hence, they may quickly become the preferred β-lactams for MABC lung disease. Finally, given the burgeoning interest in combining β-lactams from different classes to obtain synergistic effects against mycobacteria, our findings suggest several cephalosporins whose potency is enhanced by BlaMab inhibition that are worthy of combining with a fixed carbapenem/β-lactamase inhibitor combination. Cefuroxime may be of particular interest given its intrinsic potency and relatively narrow spectrum of activity. Future studies should examine the efficacy of such dual β-lactam/β-lactamase inhibitor combinations at clinically relevant exposures in preclinical models of MABC infection.

## MATERIALS AND METHODS

### Bacterial strains

*M. abscessus* strain ATCC 19977 was purchased from the American Type Culture Collection (Manassas, VA, USA) and maintained per the provider’s instructions. The MABC clinical isolates were obtained from the Johns Hopkins Hospital Clinical Microbiology Laboratory as previously described (16) and were maintained in the laboratory similarly as the ATCC 19977 strain. These isolates, from predominantly the sputum of patients with cystic fibrosis, are resistant to most antimicrobials currently available for treatment of MABC lung disease. Their drug resistance profiles were previously described (16).

### β-lactams and β-lactamase inhibitors

Faropenem, tebipenem, and biapenem were purchased from Octagon Chemicals Limited (Hangzhou, China). Doripenem hydrate, cefdinir, cefoxitin sodium salt, ceftazidime hydrate, cloxacillin sodium salt monohydrate, dicloxacillin sodium salt monohydrate, flucloxacillin, and oxacillin sodium salt were purchased from Sigma-Aldrich (St. Louis, MO, USA). Ertapenem sodium salt, cefixime trihydrate, cefpodoxime free acid, cefuroxime sodium salt, cephalexin monohydrate, cefazolin sodium salt, cefoperazone sodium salt, cefotaxime sodium salt, ceftriaxone disodium salt hemi(heptahydrate), cephalothin sodium salt, moxalactam sodium salt, aztreonam, and amoxicillin trihydrate were purchased from Research Product International (Mount Prospect, IL, USA). Imipenem monohydrate, meropenem trihydrate, and cefepime dihydrochloride monohydrate were purchased from Carbosynth (San Diego, CA, USA). Ceftaroline was manufactured by Astra-Zeneca. Vaborbactam was purchased from MedChem Express (Monmouth Junction, NJ, USA). Relebactam was purchased from Advanced ChemBlocks, Inc. (Burlingame, CA, USA). The purity of all compounds was >95%. Drug powders were stored at either 4°C or 20°C, per the manufacturers’ instructions to ensure stability. Drugs were dissolved in dimethyl sulfoxide or deionized water, again per the manufacturers’ instructions, on the day of use in MIC or disk diffusion assays.

### Media

Middlebrook 7H9 broth supplemented with 0.5% (v/v) glycerol, 10% (v/v) Middlebrook OADC enrichment, and 0.05% (v/v) Tween 80, was used as the growth medium for all routine culturing of MABC strains, including from frozen stocks. Two types of MIC assay media were used in this study. 7H9 assay medium consisted of Middlebrook 7H9 broth supplemented with 0.5% (v/v) glycerol and 10% (v/v) Middlebrook OADC enrichment, without Tween 80; CAMHB assay medium consisted only of Mueller-Hinton II broth powder. 7H11 agar supplemented with 0.5% (v/v) glycerol and 10% (v/v) Middlebrook OADC enrichment was used for disk diffusion assays. Middlebrook 7H9 broth powder, Middlebrook OADC enrichment, 7H11 agar, and CAMHB were purchased from Becton, Dickinson, and Co., (Hunt Valley, MD, USA). Glycerol and Tween 80 were purchased from Thermo Fisher Scientific (Waltham, MA, USA).

### Growth curves

MABC strains were grown from frozen stock in growth medium, shaking, at 37°C, until the cultures reached log-phase growth (optical density at 600 nm [OD_600_] between 0.4 to 0.7). The bacterial suspensions were then adjusted to an OD_600_ of 0.01 using 7H9 and CAMHB MIC assay media, in a total volume of 15 mL in 50-mL polystyrene tubes. The cultures were incubated without agitation at 30°C, and the OD_600_ of each culture was measured after 2, 3, 5, and 7 days of incubation.

### MIC broth microdilution assays

MIC assays were performed in 96-well, round-bottom plates. MABC strains were grown from frozen stock in growth medium, shaking, at 37°C, until the cultures reached log-phase growth. The bacterial suspensions were then adjusted to 1 × 10^5^ to 5 × 10^5^ colony forming units per milliliter in assay medium, and 100 μl of this bacterial suspension was added to each well, except for the no bacteria controls. Drug stock solutions prepared from powder were diluted in assay medium to achieve the appropriate concentrations in a final assay volume of 200 μl. Vaborbactam and relebactam were used at a final concentration of 4 μg/ml (31, 39), a concentration readily achievable for both β-lactamase inhibitors in human plasma (40, 41). Plates were sealed and incubated, undisturbed, at 30°C for at least 72 hours or until there was sufficient growth in the drug-free bacterial growth control wells (22). The MIC was defined as the lowest concentration of β-lactam that prevented growth as observed by the naked eye. Two biological replicates of all MIC assays were performed. MIC_50_ and MIC_90_ were defined as the MIC at which at least 50% and 90% of the clinical MABC strains were inhibited, respectively.

### Disk diffusion assays

MABC strain ATCC 19977 was grown from frozen stock to mid-log phase in growth medium until mid-log phase (OD_600_ at least 0.5). If necessary, bacterial suspensions were adjusted to an OD_600_ of 0.5, and 1 ml of the suspension was spread over 7H11 agar in a 100 mm-diameter petri dish. All drugs were prepared from powder in dimethyl sulfoxide at 10 mg/ml, further diluted in Middlebrook 7H9 broth to 1 mg/ml, and the drug solutions were applied to paper disks, 7 mm in diameter, which had been prepared from Whatman qualitative filter paper, grade 1 (Sigma-Aldrich, St. Louis, MO, USA) and sterilized prior to use. For the β-lactams, 20 μl, delivering 20 μg of drug, were applied to each disk. For the β-lactamase inhibitors alone, either 20 or 4 μl, delivering 20 or 4 μg, respectively, were applied to the disks. When used in combination with a β-lactam, 4 μl, delivering 4 μg of either relebactam or vaborbactam were applied to the appropriate disks. The disks were then air-dried and transferred using sterile forceps to the MABC-covered agar plates. The disks were pressed lightly on the agar surface to ensure contact with the bacteria. Plates were sealed in plastic bags and incubated for 4 days at 37°C.

## Acknowledgments

This study was funded by the National Institutes of Health (R21-AI137814 to ELN).

## References

1. Griffith DE, Brown-Elliott BA, Benwill JL, Wallace RJ,Jr. 2015. Mycobacterium abscessus. “Pleased to meet you, hope you guess my name…”. Ann Am Thorac Soc 12:436–439. doi: 10.1513/AnnalsATS.201501-015OI.

2. Griffith DE, Aksamit T, Brown-Elliott BA, Catanzaro A, Daley C, Gordin F, Holland SM, Horsburgh R, Huitt G, Iademarco MF, Iseman M, Olivier K, Ruoss S, von Reyn CF, Wallace RJ,Jr, Winthrop K, ATS Mycobacterial Diseases Subcommittee, American Thoracic Society, Infectious Disease Society of America. 2007. An official ATS/IDSA statement: diagnosis, treatment, and prevention of nontuberculous mycobacterial diseases. Am J Respir Crit Care Med 175:367–416. doi: 10.1164/rccm.200604-571ST.

3. Haworth CS, Banks J, Capstick T, Fisher AJ, Gorsuch T, Laurenson IF, Leitch A, Loebinger MR, Milburn HJ, Nightingale M, Ormerod P, Shingadia D, Smith D, Whitehead N, Wilson R, Floto RA. 2017. British Thoracic Society guidelines for the management of non-tuberculous mycobacterial pulmonary disease (NTM-PD). Thorax 72 Suppl 2:ii1–ii64. doi: 10.1136/thoraxjnl-2017-210927.

4. Floto RA, Olivier KN, Saiman L, Daley CL, Herrmann JL, Nick JA, Noone PG, Bilton D, Corris P, Gibson RL, Hempstead SE, Koetz K, Sabadosa KA, Sermet-Gaudelus I, Smyth AR, van Ingen J, Wallace RJ, Winthrop KL, Marshall BC, Haworth CS, US Cystic Fibrosis Foundation and European Cystic Fibrosis Society. 2016. US Cystic Fibrosis Foundation and European Cystic Fibrosis Society consensus recommendations for the management of non-tuberculous mycobacteria in individuals with cystic fibrosis. Thorax 71 Suppl 1:i1–22. doi: 10.1136/thoraxjnl-2015-207360.

5. Lee MR, Sheng WH, Hung CC, Yu CJ, Lee LN, Hsueh PR. 2015. Mycobacterium abscessus complex infections in humans. Emerg Infect Dis 21:1638–1646. doi: 10.3201/2109.141634.

6. Adjemian J, Olivier KN, Prevots DR. 2018. Epidemiology of pulmonary nontuberculous mycobacterial sputum positivity in patients with cystic fibrosis in the United States, 2010-2014. Ann Am Thorac Soc 15:817–826. doi: 10.1513/AnnalsATS.201709-7270C.

7. Mougari F, Guglielmetti L, Raskine L, Sermet-Gaudelus I, Veziris N, Cambau E. 2016. Infections caused by Mycobacterium abscessus: epidemiology, diagnostic tools and treatment. Expert Rev Anti Infect Ther 14:1139–1154. doi: 10.1080/14787210.2016.1238304.

8. Rominski A, Schulthess B, Müller DM, Keller PM, Sander P. 2017. Effect of β-lactamase production and β-lactam instability on MIC testing results for Mycobacterium abscessus. J Antimicrob Chemother 72:3070–3078. doi: 10.1093/jac/dkx284.

9. Soroka D, Dubée V, Soulier-Escrihuela O, Cuinet G, Hugonnet JE, Gutmann L, Mainardi JL, Arthur M. 2014. Characterization of broad-spectrum Mycobacterium abscessus class A β-lactamase. J Antimicrob Chemother 69:691–696. doi: 10.1093/jac/dkt410.

10. Kwon HH, Tomioka H, Saito H. 1995. Distribution and characterization of β-lactamases of mycobacteria and related organisms. Tuber Lung Dis 76:141–148. doi: 10.1016/0962- 8479(95)90557-X.

11. Fattorini L, Oliva B, Orefici G. 1986. Expression and some properties of β-lactamase from Mycobacterium fortuitum. Drugs Exp Clin Res 12:973–977.

12. Zhang Y, Steingrube VA, Wallace RJ,Jr. 1992. β-Lactamase inhibitors and the inducibility of the β-lactamase of Mycobacterium tuberculosis. Am Rev Respir Dis 145:657–660. doi: 10.1164/ajrccm/145.3.657.

13. Ramírez A, Ruggiero M, Aranaga C, Cataldi A, Gutkind G, de Waard JH, Araque M, Power P. 2017. Biochemical characterization of β-lactamases from Mycobacterium abscessus complex and genetic environment of the β-lactamase-encoding gene. Microb Drug Resist 23:294–300. doi: 10.1089/mdr.2016.0047.

14. Kaushik A, Makkar N, Pandey P, Parrish N, Singh U, Lamichhane G. 2015. Carbapenems and rifampin exhibit synergy against Mycobacterium tuberculosis and Mycobacterium abscessus. Antimicrob Agents Chemother 59:6561–6567. doi: 10.1128/AAC.01158-15.

15. Soroka D, Ourghanlian C, Compain F, Fichini M, Dubée V, Mainardi JL, Hugonnet JE, Arthur M. 2017. Inhibition of β-lactamases of mycobacteria by avibactam and clavulanate. J Antimicrob Chemother 72:1081–1088. doi: 10.1093/jac/dkw546.

16. Kaushik A, Gupta C, Fisher S, Story-Roller E, Galanis C, Parrish N, Lamichhane G. 2017. Combinations of avibactam and carbapenems exhibit enhanced potencies against drug-resistant Mycobacterium abscessus. Future Microbiol 12:473–480. doi: 10.2217/fmb-2016-0234.

17. Lefebvre AL, Le Moigne V, Bernut A, Veckerlé C, Compain F, Herrmann JL, Kremer L, Arthur M, Mainardi JL. 2017. Inhibition of the β-lactamase BlaMab by avibactam improves the in vitro and in vivo efficacy of imipenem against Mycobacterium abscessus. Antimicrob Agents Chemother 61:e02440–16. doi: 10.1128/AAC.02440-16.

18. Dubée V, Bernut A, Cortes M, Lesne T, Dorchene D, Lefebvre AL, Hugonnet JE, Gutmann L, Mainardi JL, Herrmann JL, Gaillard JL, Kremer L, Arthur M. 2015. β-lactamase inhibition by avibactam in Mycobacterium abscessus. J Antimicrob Chemother 70:1051–1058. doi: 10.1093/jac/dku510.

19. Dubée V, Soroka D, Cortes M, Lefebvre AL, Gutmann L, Hugonnet JE, Arthur M, Mainardi JL. 2015. Impact of β-lactamase inhibition on the activity of ceftaroline against Mycobacterium tuberculosis and Mycobacterium abscessus. Antimicrob Agents Chemother 59:2938–2941. doi: 10.1128/AAC.05080-14.

20. Lefebvre AL, Dubée V, Cortes M, Dorchêne D, Arthur M, Mainardi JL. 2016. Bactericidal and intracellular activity of β-lactams against Mycobacterium abscessus. J Antimicrob Chemother 71:1556–1563. doi: 10.1093/jac/dkw022.

21. Zhanel GG, Lawrence CK, Adam H, Schweizer F, Zelenitsky S, Zhanel M, Lagacé-Wiens PRS, Walkty A, Denisuik A, Golden A, Gin AS, Hoban DJ, Lynch JP,3rd, Karlowsky JA. 2018. Imipenem-relebactam and meropenem-vaborbactam: Two novel carbapenem-β-lactamase inhibitor combinations. Drugs 78:65–98. doi: 10.1007/s40265-017-0851-9.

22. Clinical and Laboratory Standards Institute (CLSI). 2011. Susceptibility testing of mycobacteria, nocardiae, and other aerobic actinomycetes; Approved standard - Second edition. CLSI document M24-A2. Vol. 31 No. 5. Clinical and Laboratory Standards Institute, Wayne, PA, USA.

23. Viaene E, Chanteux H, Servais H, Mingeot-Leclercq MP, Tulkens PM. 2002. Comparative stability studies of antipseudomonal β-lactams for potential administration through portable elastomeric pumps (home therapy for cystic fibrosis patients) and motor-operated syringes (intensive care units). Antimicrob Agents Chemother 46:2327–2332. doi: 10.1128/AAC.46.8.2327-2332.2002.

24. Lemaire S, Van Bambeke F, Mingeot-Leclercq MP, Tulkens PM. 2005. Activity of three β-lactams (ertapenem, meropenem and ampicillin) against intraphagocytic Listeria monocytogenes and Staphylococcus aureus. J Antimicrob Chemother 55:897–904. doi: 10.1093/jac/dki094.

25. Srivastava S, van Rijn SP, Wessels AM, Alffenaar JW, Gumbo T. 2016. Susceptibility testing of antibiotics that degrade faster than the doubling time of slow-growing mycobacteria: Ertapenem sterilizing effect versus Mycobacterium tuberculosis. Antimicrob Agents Chemother 60:3193–3195. doi: 10.1128/AAC.02924-15.

26. Cortes MA, Nessar R, Singh AK. 2010. Laboratory maintenance of Mycobacterium abscessus. Curr Protoc Microbiol Chapter 10:Unit 10D.1. doi: 10.1002/9780471729259.mc10d01s18.

27. BD Diagnostics-Diagnostic Systems. 2009. Difco & BBL Manual. Manual of Microbiological Culture Media. Becton, Dickinson and Company, Sparks, MD, USA. https://www.bd.com/resource.aspx?IDX=9572.

28. Larsen MH, Biermann K, Jacobs WR,Jr. 2007. Laboratory maintenance of Mycobacterium tuberculosis. Curr Protoc Microbiol Chapter 10:Unit 10A.1. doi: 10.1002/9780471729259.mc10a01s6.

29. Ripoll F, Pasek S, Schenowitz C, Dossat C, Barbe V, Rottman M, Macheras E, Heym B, Herrmann JL, Daffé M, Brosch R, Risler JL, Gaillard JL. 2009. Non mycobacterial virulence genes in the genome of the emerging pathogen Mycobacterium abscessus. PLoS One 4:e5660. doi: 10.1371/journal.pone.0005660.

30. Kumar P, Chauhan V, Silva JRA, Lameira J, d’Andrea FB, Li SG, Ginell SL, Freundlich JS, Alves CN, Bailey S, Cohen KA, Lamichhane G. 2017. Mycobacterium abscessus L,D-transpeptidases are susceptible to inactivation by carbapenems and cephalosporins but not penicillins. Antimicrob Agents Chemother 61:e00866-17. doi: 10.1128/AAC.00866-17.

31. Clinical and Laboratory Standards Institute (CLSI). 2017. Performance standards for antimicrobial susceptibility testing, M100, 27th edition. Clinical and Laboratory Standards Institute, Wayne, PA, USA.

32. Moine P, Fish DN. 2013. Pharmacodynamic modelling of intravenous antibiotic prophylaxis in elective colorectal surgery. Int J Antimicrob Agents 41:167–173. doi: 10.1016/j.ijantimicag.2012.09.017.

33. Story-Roller E, Lamichhane G. 2018. Overcoming β-lactam resistance in Mycobacterium abscessus. Abstract 803, IDWeek 2018 conference, 2-6 October 2018, San Francisco, California, USA. https://idsa.confex.com/idsa/2018/webprogram/start.html.

34. Beganovic M, Luther MK, Rice LB, Arias CA, Rybak MJ, LaPlante KL. 2018. A review of combination antimicrobial therapy for Enterococcus faecalis bloodstream infections and infective endocarditis. Clin Infect Dis 67:303–309. doi: 10.1093/cid/ciy064.

35. Gonzalo X, Drobniewski F. 2013. Is there a place for β-lactams in the treatment of multidrug-resistant/extensively drug-resistant tuberculosis? Synergy between meropenem and amoxicillin/clavulanate. J Antimicrob Chemother 68:366–369. doi: 10.1093/jac/dks395.

36. Story-Roller E, Maggioncalda EC, Cohen KA, Lamichhane G. 2018. Mycobacterium abscessus and β-lactams: Emerging insights and potential opportunities. Front Microbiol 9:2273. doi: 10.3389/fmicb.2018.02273.

37. Lavollay M, Fourgeaud M, Herrmann JL, Dubost L, Marie A, Gutmann L, Arthur M, Mainardi JL. 2011. The peptidoglycan of Mycobacterium abscessus is predominantly cross-linked by L,D-transpeptidases. J Bacteriol 193:778–782. doi: 10.1128/JB.00606-10.

38. Pandey R, Chen L, Shashkina E, Manca C, Bonomo RA, Jenkins SG, Kreiswirth BN. 2017. Evaluation of ceftaroline-avibactam activity in vitro and ex vivo against Mycobacterium abscessus complex. Abstract 2276, IDWeek 2017 conference, 4-8 October 2017, San Diego, California, USA. https://idsa.confex.com/idsa/2017/webprogram/start.html.

39. Lomovskaya O, Sun D, Rubio-Aparicio D, Nelson K, Tsivkovski R, Griffith DC, Dudley MN. 2017. Vaborbactam: Spectrum of β-lactamase inhibition and impact of resistance mechanisms on activity in Enterobacteriaceae. Antimicrob Agents Chemother 61: e01443–17. doi: 10.1128/AAC.01443-17.

40. Rhee EG, Rizk ML, Calder N, Nefliu M, Warrington SJ, Schwartz MS, Mangin E, Boundy K, Bhagunde P, Colon-Gonzalez F, Jumes P, Liu Y, Butterton JR. 2018. Pharmacokinetics, safety, and tolerability of single and multiple doses of relebactam, a β-lactamase inhibitor, in combination with imipenem and cilastatin in healthy participants. Antimicrob Agents Chemother 62:10.1128/AAC.00280-18. Print 2018 Sep. doi: 10.1128/AAC.00280-18.

41. Griffith DC, Loutit JS, Morgan EE, Durso S, Dudley MN. 2016. Phase 1 study of the safety, tolerability, and pharmacokinetics of the β-lactamase inhibitor vaborbactam (RPX7009) in healthy adult subjects. Antimicrob Agents Chemother 60:6326–6332. doi: 10.1128/AAC.00568-16.

